# Hippocampal astroglial hypertrophy in mice subjected to early life maternal deprivation

**DOI:** 10.1101/2021.09.05.459015

**Authors:** Arnab Nandi, Garima Virmani, Swananda Marathe

## Abstract

Early-life stress (ELS), including chronic deprivation of maternal care, exerts persistent life-long effects on animal physiology and behavior, and is associated with several neurodevelopmental disorders. Long-lasting changes in neuronal plasticity and electrophysiology are documented extensively in the animal models of ELS. However, the role of astroglia in the lasting effects of ELS remains elusive. Astrocytes are intricately involved in the regulation of synaptic physiology and behavior. Moreover, astrocytes play a major role in the innate and adaptive immune responses in the central nervous system (CNS). The role of immune responses and neuroinflammation in the altered brain development and persistent adverse effects of ELS are beginning to be explored. Innate immune response in the CNS is characterized by a phenomenon called astrogliosis, a process in which astrocytes undergo hypertrophy, along with changes in gene expression and function. While the immune activation and neuroinflammatory changes concomitant with ELS, or in juveniles and young adults have been reported, it is unclear whether mice subjected to ELS exhibit astrogliosis-like alterations well into late-adulthood. Here, we subjected mice to maternal separation from postnatal day 2 to day 22 and performed comprehensive morphometric analysis of hippocampal astrocytes during late-adulthood. We found that the astrocytes in the *stratum radiatum* region of the CA1 hippocampal subfield from maternally separated mice exhibit significant hypertrophy as late as 8 months of age, revealing the crucial changes in astrocytes that manifest long after the cessation of ELS. This study highlights the persistence of neuroinflammatory changes in mice exposed to ELS.

## Introduction

Chronic early-life stress (ELS) has detrimental effects on neonatal brain development resulting in impaired adult brain function (Smith and Pollak, 2020). As a result, ELS acts as a major risk factor for neurodevelopmental disorders such as schizophrenia, affective disorders and learning disabilities (Akdeniz et al., 2014; Schmitt et al., 2014; Syed and Nemeroff, 2017; Hambrick et al., 2019; Smith and Pollak, 2020). Investigating the molecular and cellular processes underlying the correlation between early-life adversity and neurodevelopmental disorders would help in designing targeted therapies for such disorders. Although several neuronal and synaptic mechanisms have been proposed (Chen and Baram, 2015; Lopatina et al., 2021), glial cells have received relatively less scrutiny.

Glial cells, especially astrocytes, have been shown to play a major role in the regulation of synaptic transmission and behavior (Ota et al., 2013; De Pittà et al., 2016; Nicola J. Allen, 2017; Santello et al., 2019; Perez-Catalan et al., 2021). Interestingly, glial development is temporally delayed as compared to neuronal development, and a large part of it coincides with the stressful experiences of early life (Sauvageot and Stiles, 2002; Molofsky and Deneen, 2015). Hence, it is tempting to speculate that the neonatal glial development is impaired as a result of ELS, and this underlies the pathophysiology of the resulting neurodevelopmental disorders. Moreover, glial cells also play a major role in the innate and adaptive immune responses in the central nervous system (CNS) (Carpentier et al., 2005; Colombo and Farina, 2016). Heightened neuro-immune responses and neuroinflammation have been implicated in the detrimental effects of ELS (Ader and Sb, 1965; Solomon et al., 1968; Shanks and Lightman, 2001; Hennessy et al., 2010; Ganguly and Brenhouse, 2015; Danese and J Lewis, 2016). Additionally, immune challenges in early life have been shown to replicate many of the adverse effects of ELS (Bilbo and Schwarz, 2009; Patterson, 2009; Knuesel et al., 2014; Danese and J Lewis, 2016; Cao et al., 2021). Given the crucial role played by astrocytes in neuroimmune responses and neuroinflammation (Cekanaviciute and Buckwalter, 2016; Colombo and Farina, 2016; Liddelow and Barres, 2017), astrocytes may prove to be the prime culprits in the deleterious effects of ELS.

In response to an injury or a trauma to the CNS, astrocytes undergo a phenomenon called reactivation, which is characterized by profound changes to their structure, gene expression, physiology and function (Barres, 2008; Sofroniew, 2009, 2014; Molofsky et al., 2012; Burda and Sofroniew, 2014; Chung et al., 2015). The process of astroglial reactivation is believed to be consequential to the disease outcome, and the neuroprotective astrocytes have been hypothesized to hold enormous therapeutic potential (Liddelow and Barres, 2017; Escartin et al., 2021). Marked hypertrophy of astroglia is a hallmark of reactivation, which is an integral part of neuroinflammatory machinery (Sofroniew, 2009; Burda and Sofroniew, 2014; Chung et al., 2015; Cekanaviciute and Buckwalter, 2016; Colombo and Farina, 2016; Liddelow and Barres, 2017). Hence, the study of astroglial structure is an important first step in assessing neuroimmune function. Moreover, hippocampal astroglial hypertrophy is associated with normal ageing and is typically seen from 14-18 months of age in rodents (Rodríguez et al., 2014; Bondi et al., 2021). Here we asked if hippocampal astroglia exhibit changes in structure during late-adulthood (8 months of age) after exposure to ELS in mice.

Astrocytic structure is highly dynamic and astrocytes show plasticity in response to physiological and pathological stimuli (Schiweck et al., 2018). Interestingly, hippocampal astrocytes exhibit structural plasticity following chronic stress and chronic antidepressant therapy (Sethi et al., 2021; Virmani et al., 2021). Astrocytes have also been shown to undergo alterations in maternally separated (MS) animals, although the results have been inconsistent. Glial Fibrillary Acidic Protein (GFAP) has been used to assess the astroglial changes in response to ELS. In rats subjected to ELS, reduced GFAP immunoreactivity and morphological complexity was shown in male pups (Roque et al., 2016; Saavedra et al., 2017), Another study observed no differences in the density of hippocampal astrocytes on postnatal day 14 (Musholt et al., 2009). However, at 2 months of age, GFAP mRNA expression (Burke et al., 2013), and the number and morphology of GFAP-positive astrocytes was unchanged (Gondré-Lewis et al., 2016). On the other hand, the density of GFAP-positive astrocytes was decreased in 10 month old MS rats (Leventopoulos et al., 2007). In summary, the changes to GFAP-positive hippocampal astrocytes as a result of ELS have not led to any clear conclusions. Moreover, for these changes to be responsible for the persistent adverse effects of ELS, astrocytes need to be studied at late time points. Given the complex morphology that astrocytes assume *in vivo*, a comprehensive study of astrocyte morphometry needs to be undertaken to tease apart the changes to astrocyte structure.

Here, we sought to address the changes in astrocytic structure in the hippocampal astrocytes during late adulthood in MS mice. We separated the pups from their mothers for 3 hours each day from the postnatal day 2 (P2) to postnatal day 22 (P22). The time-window for MS was chosen so as to cover the postnatal astrogliogenesis in developing mice. We found significant hypertrophy in the astrocytes in 8 month old MS mice in hippocampal CA1 *stratum radiatum*, an important output subfield of the hippocampus. These results demonstrate long-lasting detrimental effects of ELS on hippocampal astroglia, which may have important implications to basal astroglial functions and their immune responses.

## Materials and Methods

### Animals

C57BL/6J mice used in this study were bred and housed in the IISc Central Animal Facility. Mice were maintained on a 12/12 hour light/dark cycle and had access to food and water *ad libitum*. Experiments were carried out in accordance with the protocols approved by the Institutional Animal Ethics Committee (IAEC). All efforts were made to reduce the number of animals by using the principle of 3R’s.

### Maternal separation (MS) paradigm

Mouse pups were separated from the dams from P2 to P22 for 3 hours each day. Dams were carefully removed from the home cages and were kept in a fresh cage, while the pups remained in the home cages during separation. The cages with pups were kept on a heating pad maintained at 32-34°C. After 3 hours of MS, the dams were returned to their respective home cages. Control dams were handled similarly, but without being separated from their pups. All mice were left undisturbed from P23 onwards. The pups were weaned from their mothers at P30, and were kept undisturbed until 8 months of age.

### Tissue preparation

8 month old male mice were anesthetized using isoflurane and were transcardially perfused with 0.9% saline followed by 4% paraformaldehyde (PFA) prepared in 0.1M phosphate buffer (PB). Brains were dissected out and post-fixed overnight in 4% PFA. Following this, the brains were cryoprotected in a 30% sucrose solution made in 0.1M PB. Brains were then frozen in the blocks of OCT compound (Sigma), and stored at −80°C until sectioning. 40µm coronal sections were cut on a cryostat (Leica, CM 1850) and were stored at 4°C in 0.1 M PB until staining.

### Immunofluorescence and imaging

Immunofluorescence was performed as described previously (Virmani et al., 2021). In brief, sections were washed thrice in 0.1M PB and followed by a 1-2 hour incubation with a blocking solution (1% normal donkey serum, 3% bovine serum albumin, 0.3% Triton X-100, prepared in 0.1 M PB) at room temperature. This was followed by an overnight incubation with the primary antibody at 4°C (chicken pAb anti-GFAP, 1:1000; Novus, NBP1-05198). Sections were again washed thrice in 0.1 M PB and then incubated with the secondary antibody for 2 h (goat anti-chicken-IgY conjugated to Alexa Fluor 594, Abcam, ab150172). Following the incubation with the secondary antibody, sections were washed again with 0.1 M PB four times. Sections were mounted on glass slides in the mounting medium containing DAPI (Abcam, ab104139). The slides were imaged on a Zeiss LSM 880 airyscan confocal microscope with a Plan-Apochromat 20X/0.8NA M27 objective. 1024 × 1024 pixels, 8-bit images were acquired with 1 μm z-step size. Image metadata was used for pixels to micrometers conversion.

### Image analysis

Images of single astrocytes used for morphological analysis were generated using a custom macro in ImageJ. In brief, maximum intensity projection of confocal images were generated. The 2-D images, thus generated, were then binarized. We then used the “analyze particles” tool to eliminate the background using size exclusion. Following this, we used morphological closing on binarized images. Next, we filled holes in the thresholded image using the “fill holes” tool. Following this step, the cells were skeletonized using the “skeletonize” plugin to generate cells with single pixel-wide processes. We then ran the “analyze particles” tool again to generate ROI for each cell skeleton. After this, we cropped the individual cells by selecting from the ROI manager in ImageJ in contrast-enhanced maximum intensity projection images. Contrast enhancement was done to capture the cell morphology well and it was done for all the images in a similar manner. These cropped cells were then used for morphological feature extraction. Cropped cells from CA1 *stratum radiatum* were analyzed using the SMorph software as described earlier (Sethi et al., 2021).

### Statistical analysis

All statistical analyses were done using GraphPad Prism 8.0.2 and the results were expressed as box and whiskers plots for 2 group comparisons, and mean ± SEM for Sholl analysis. Boxes denote the 25th to 75th percentile and whiskers denote the minimum and maximum values in the datasets. Median is represented as a horizontal line across the box and the mean is represented as “+”. For 2 group comparisons, normality of the data was determined using Kolmogorov-Smirnov normality test with Dallal-Wilkinson-Lilliefor approximation, with the distribution considered normal (Gaussian) at the alpha of 0.05. Comparisons were made using student’s *t*-test for normally distributed data and using Mann-Whitney *U*-test for the data that was not normally distributed. Sholl analysis data was analyzed using *t*-test with false discovery rate to correct for multiple comparisons. The differences between the 2 groups was deemed to be significant at p < 0.05.

## Results

### Analysis of the size of hippocampal astrocytes in maternally separated mice

To assess whether ELS affects the cell size of hippocampal astrocytes, pups were separated from the dams for 3 hours each day from P2 until P22. Control mice on the other hand were handled similarly, but were never separated from the dams. Since astrocytes have a complex stellate morphology, we computed multiple parameters related to cell size; namely, 2-D projected surface area, area of the convex hull, the maximum radius of cells and the total length of the cell skeleton. We found a statistically significant 39% increase in the 2-D projected area of GFAP-positive profiles of astrocytes (Control: 184.7 ± 8.25 µm^2^, MS: 257 ± 13.03 µm^2^, p<0.0001, Mann-Whitney *U*-test) (Figure 1A). We further analysed the area of the convex hull drawn around the extremities of the cell skeleton. We found a statistically significant 47% increase in the area of convex hull in MS mice as compared to the controls (Control: 379.6 ± 19.71 µm^2^, MS: 558.6 ± 30.29 µm^2^, p<0.0001, Mann-Whitney *U*-test) (Figure 1B). We next computed the maximum radius as the distance from the center of the soma to the furthermost point on the cell. We found a statistically significant 24% increase in the maximum radius of the cells in MS mice compared to the controls (Control: 16.99 ± 0.4738 µm, MS: 21.13 ± 0.6614 µm, p<0.0001, Mann-Whitney *U*-test) (Figure 1C). We further analyzed the total length of the cell skeleton in MS mice and compared it to controls. We found that the MS mice showed a statistically significant 36% increase in the skeleton length as compared to controls (Control: 84.63 ± 3.551 µm, MS: 115.5 ± 5.308 µm, p<0.0001, Mann-Whitney *U*-test) (Figure 1D). Taken together, these results reveal a robust increase in cell size of the hippocampal astrocytes in MS mice as compared to the controls.

**Figure 1:**
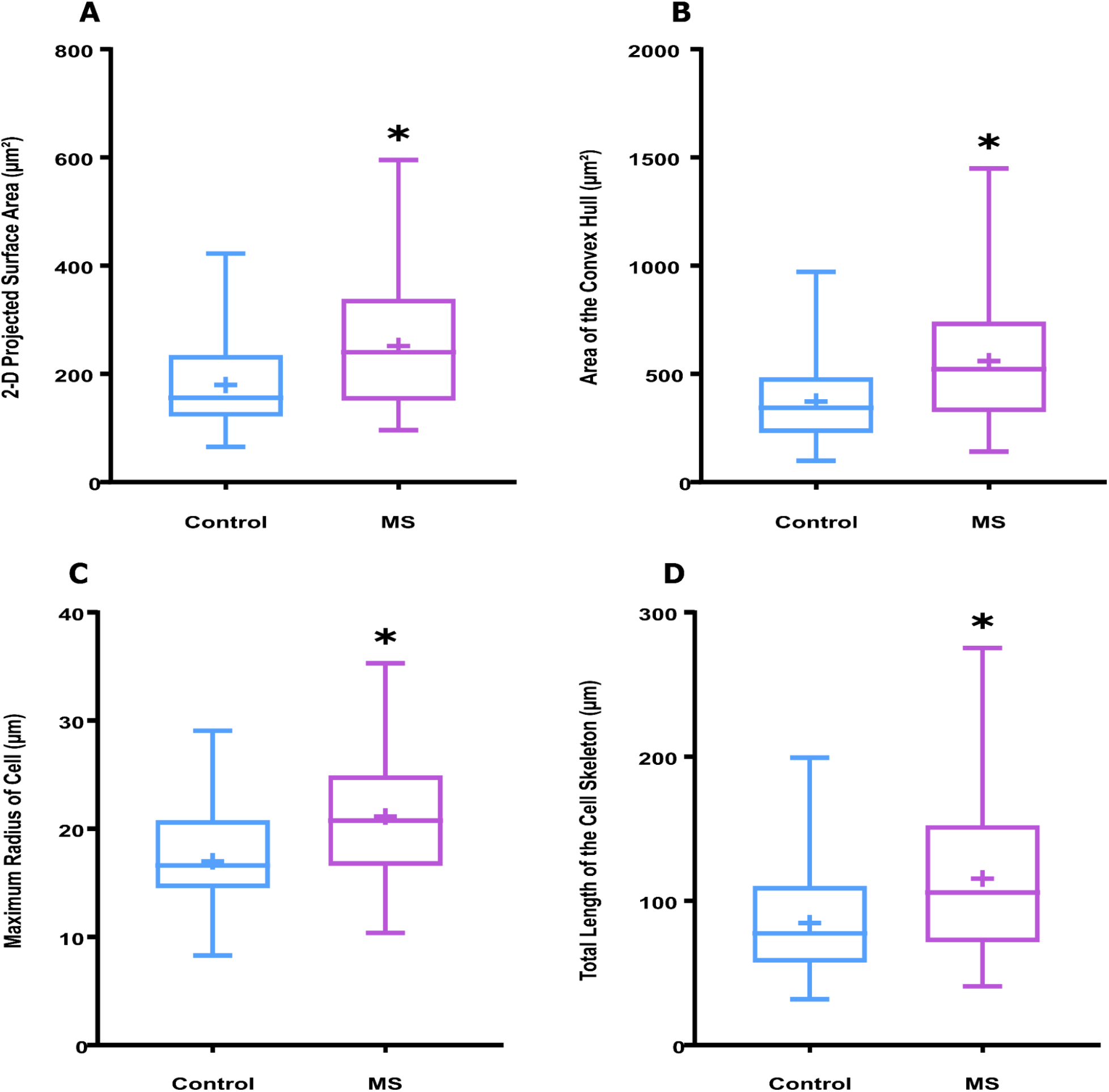
Analysis of the size of hippocampal astrocytes in maternally separated mice. We assessed the parameters related to the cell size of hippocampal astrocytes in MS animals and controls. Box and whiskers plots show a statistically significant increase in the **(A)** 2D-projected surface area, **(B)** area of the convex hull, **(C)** maximum radius and **(D)** the total skeleton length. n= 90-100 cells per group sampled evenly from 4 mice per group. All comparisons made using Mann-Whitney *U*-test. * represents p<0.05.

### Analysis of the astrocytic arbors in the hippocampus of maternally separated mice

In order to study the changes in astrocytic branches as a result of ELS (Figure 2A), we analysed the number of different branch classes and branch points. The analysis involved the number of primary, secondary, tertiary and terminal branches and the number of branch points. Interestingly, we found no difference in the number of primary processes between the control mice and MS-treated mice (Control: 4.282 ± 0.1341, MS: 4.225 ± 0.1697, p=0.5577, Mann-Whitney *U*-test) (Figure 2B). On the other hand, we found a statistically significant increase in the number of secondary and tertiary branches in the MS-treated mice compared to the controls (Secondary branches: Control: 5.544 ± 0.2734, MS: 7 ± 0.2542, p<0.0001, Mann-Whitney *U*-test) (Tertiary branches: Control: 4.845 ± 0.3223, MS: 6.955 ± 0.3871, p<0.0001, Mann-Whitney *U*-test) (Figure 2C, 2D). We further analyzed the number of terminal branches per cell and found a statistically significant increase in the number of terminal branches in the MS mice as compared to controls (Control: 12.82 ± 0.5138, MS: 17.48 ± 0.7545, p<0.0001, Mann-Whitney *U*-test) (Figure 2E). We also compared the number of branch points between the controls and MS groups and found a significant increase in the MS mice compared to controls (Control: 7.66 ± 0.3649, MS: 11.33 ± 0.5663, p<0.0001, Mann-Whitney *U*-test) (Figure 2F).

**Figure 2:**
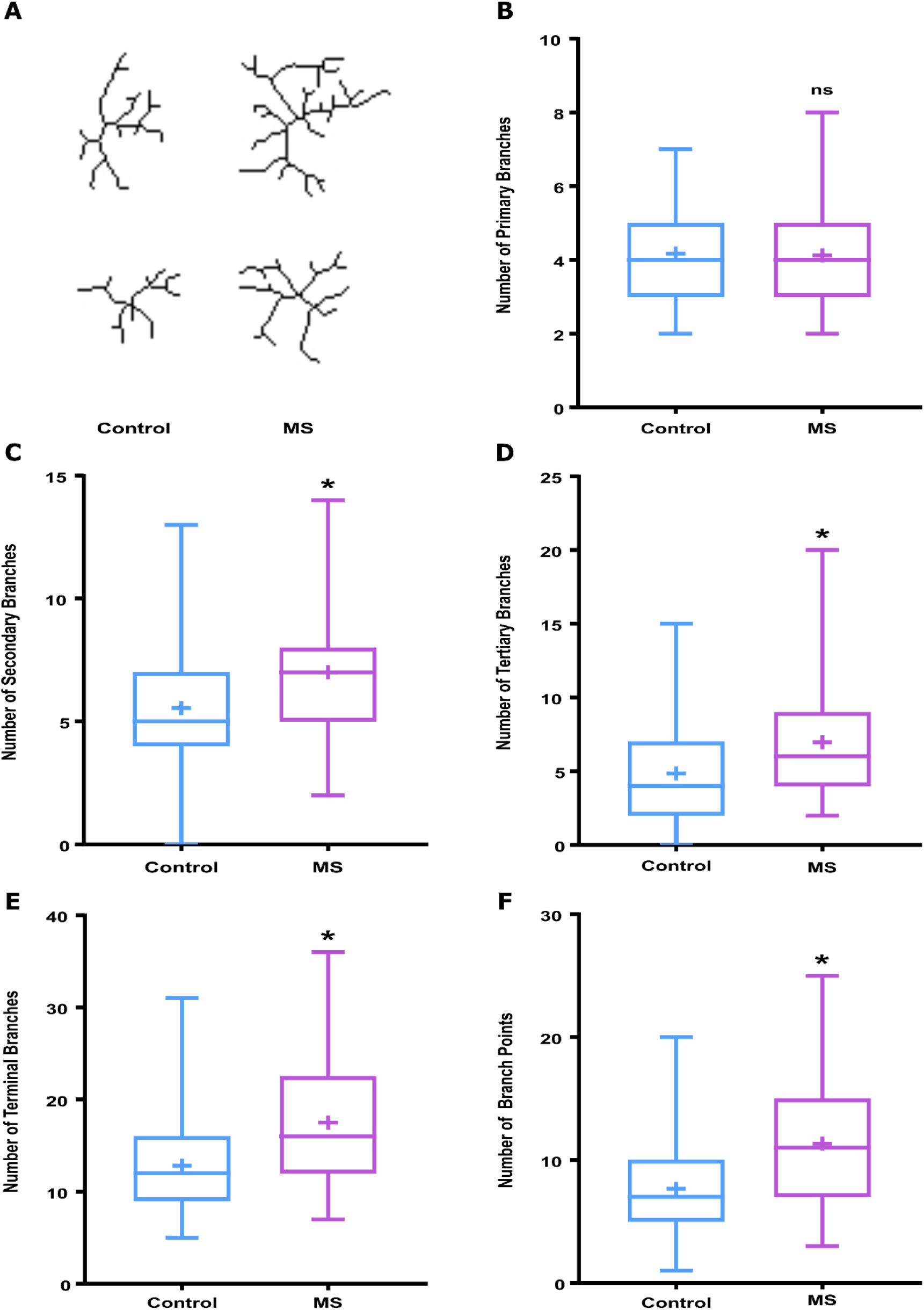
Analysis of the astrocytic arbors in the hippocampus of maternally separated mice. We analyzed the parameters related to the branching of astrocytic processes in controls and MS-treated mice. **(A)** Representative skeletons of cells belonging to control mice and MS mice are shown. **(B)** Box and whiskers plot shows no statistically significant change between the two groups in the number of primary branches. Box and whiskers plots show a statistically significant increase in the number of **(C)** secondary, **(D)** tertiary, and **(E)** terminal branches as well as **(F)** the number of branch points in MS mice as compared to the controls. n= 90-100 cells per group sampled evenly from 4 mice per group. All comparisons made using Mann-Whitney *U*-test. * represents p<0.05.

We also analysed the average lengths of the primary, secondary and tertiary branches. We found no statistically significant differences in the average lengths of primary (Control: 4.775 ± 0.1258 µm, MS: 4.744 ± 0.1454 µm, p=0.6836, Mann-Whitney *U*-test) (Figure S1A), secondary (Control: 4.194 ± 0.1254 µm, MS: 4.259 ± 0.1128 µm, p=0.9384, Mann-Whitney *U*-test) (Figure S1B) or tertiary (Control: 3.499 ± 0.1772 µm, MS: 3.598 ± 0.123 µm, p=0.658, Student’s *t*-test) (Figure S1C) branches between the controls and MS mice.

### Sholl analysis of hippocampal astrocytes in maternally separated mice

Furthermore, to study the effects of ELS on astrocyte arborization, we performed Sholl analysis on the processes of hippocampal astrocytes. We found a significant increase in the ramification of astrocytic processes in the MS mice as compared to the controls (Figure 3A), along with a statistically significant increase in the area under the Sholl analysis curve (Control: 45.68 ± 1.847 µm, MS: 62.26 ± 2.902 µm, p<0.0001, Mann-Whitney *U*-test) (Figure 3B). We next computed the critical radius, the radius corresponding to the peak in the Sholl analysis curve. We found a statistically significant increase in the critical radius in MS animals as compared to the controls (Control: 7.859 ± 0.3197 µm, MS: 9.632 ± 0.3467 µm, p=0.0002, Mann-Whitney *U*-test) (Figure 3C). On the other hand, the critical value, corresponding to the height of the peak of the Sholl analysis curve, did not change as a result of MS (Control: 5.59 ± 0.1616 µm, MS: 6.202 ± 0.2231 µm, p=0.0611, Mann-Whitney U-test) (Figure 3D). Taken together, these results show that MS induces robust hypertrophy in the hippocampal astrocytes at 8 months of age.

**Figure 3:**
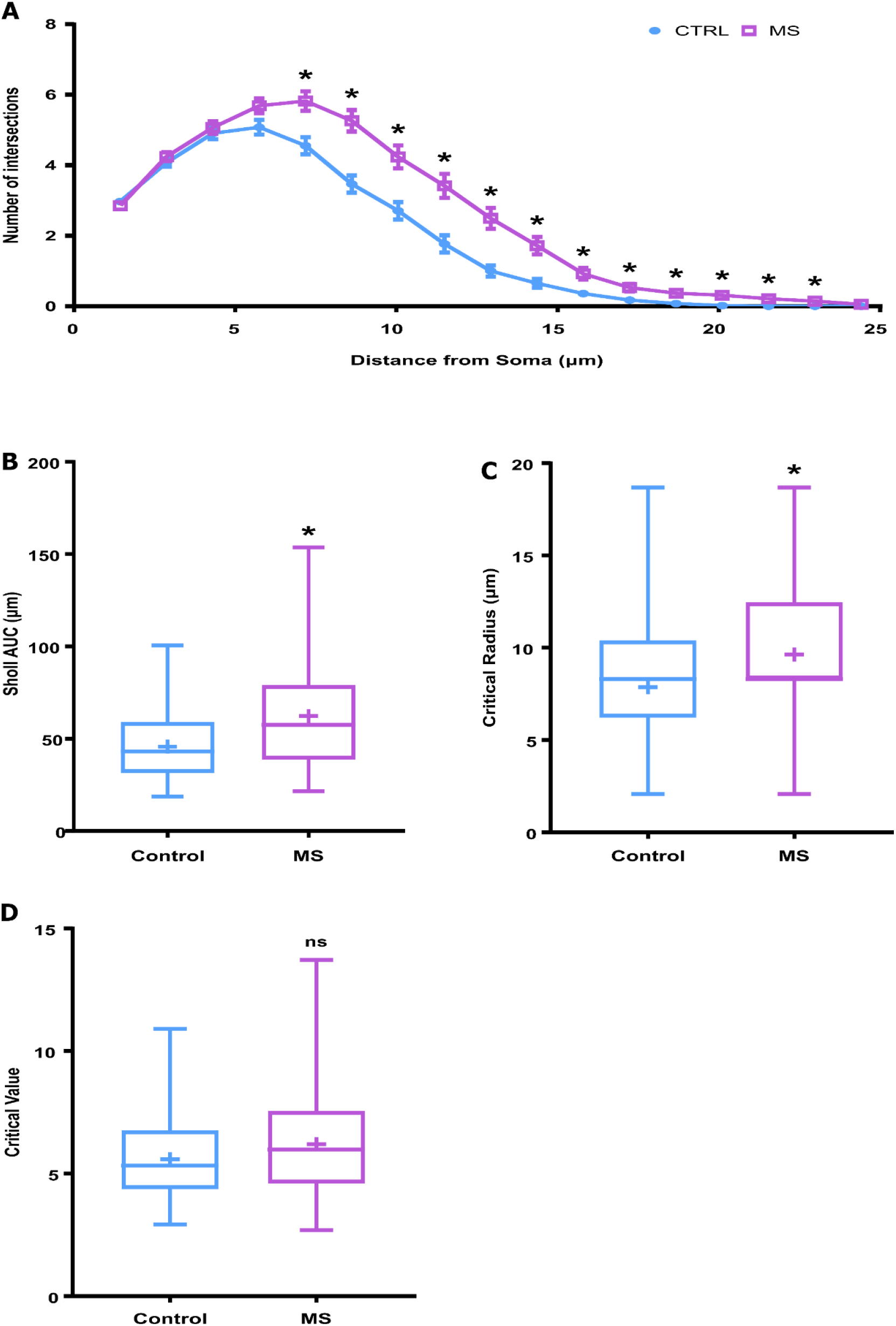
Sholl analysis of hippocampal astrocytes in maternally separated mice. We analyzed astrocyte arborization using Sholl analysis and compared the process ramifications between the MS animals and controls. **(A)** Line plot shows a statistically significant increase in the arborization in astrocytes of MS animals as compared to the controls. Box and whiskers plot shows a statistically significant increase in **(B)** the area under the Sholl analysis curve and **(C)** the critical radius in MS mice compared to controls. However, **(D)** the critical value shows no statistically significant difference between the two groups. n= 90-100 cells per group sampled evenly from 4 mice per group. Sholl analysis results in (A) were analysed using multiple *t-tests* with false discovery rate (FDR) approach to correct for multiple comparisons. All other comparisons were made using Mann-Whitney *U*-test. * represents p<0.05.

## Discussion

Astrocytes have historically been thought to play a supportive role in the CNS. However, emerging evidence highlights their role in directly modulating synaptic responses and behavior (De Pittà et al., 2016; Santello et al., 2019). Astrocytes largely develop postnatally in the rodent brain and these developmental processes temporally overlap with the timeframes typically used in the ELS paradigms (Molofsky and Deneen, 2015; Hambrick et al., 2019). We speculate that the MS paradigm used in this study affects astroglial development and that the astrocytes play a crucial role in the manifestation of the long-lasting impairments caused by ELS. Given that ELS in humans is known to be a major risk factor for several neurodevelopmental disorders (Akdeniz et al., 2014; Schmitt et al., 2014; Syed and Nemeroff, 2017; Hambrick et al., 2019; Smith and Pollak, 2020), it is crucial to investigate the underlying molecular and cellular mechanisms.

Astrocytes possess highly branched stellate morphology, which is highly plastic (Bernardinelli et al., 2014; Schiweck et al., 2018). The fine leaflet-like processes dynamically contact synapses forming perisynaptic astrocytic processes (PAPs) (Ghézali et al., 2016; Schiweck et al., 2018). PAPs show activity-dependent motility and play an integral role in synaptic plasticity (Bernardinelli et al., 2014). Furthermore, astrocytes can sense the release of neurotransmitters in the synaptic cleft through PAPs and release gliotransmitters in response, which profoundly modulate pre- and post-synaptic responses (Harada et al., 2016; Savtchouk and Volterra, 2018). Along with these basal functions, astrocytes also play an integral role in innate and adaptive immune responses in the CNS including neuroinflammatory pathways (Carpentier et al., 2005; Cekanaviciute and Buckwalter, 2016). Astrocytes show context-dependent dichotomous effects that either exacerbate or alleviate the tissue damage (Colombo and Farina, 2016). In agreement with this dichotomy, reactive astrocytes are proposed to polarize into neurotoxic (A1) and neuroprotective (A2) phenotypes, resulting in highly divergent outcomes (Liddelow and Barres, 2017; Escartin et al., 2021). A distinguishing feature of reactive astrocytes, irrespective of their underlying molecular and functional phenotypes, is prominent morphological hypertrophy. Here, we investigated whether the hippocampal astrocytes show hypertrophy during late-adulthood in mice subjected to neonatal MS.

We found that the hippocampal astrocytes in the 8 month old MS mice show hypertrophic morphology as compared to the control mice. This hypertrophy is manifested both in terms of an increase in the cell size and the complexity of arbors. Although changes to GFAP-positive astrocytes had been studied in pups and very young adults (Musholt et al., 2009; Burke et al., 2013; Gondré-Lewis et al., 2016), the changes we observe reveal that the effects of ELS on astrocytes are long-lasting, and hence may underlie the persistent adverse effects of ELS. Furthermore, to our knowledge, this is the first report showing astroglial hypertrophy in 8 month old maternally separated mice, using comprehensive morphometric analysis of GFAP-positive profiles.

The hypertrophy seen in the hippocampal astrocytes of maternally separated mice strongly hints at a possible reactive phenotype. Although astrocytes show a high degree of structural plasticity (Schiweck et al., 2018; Sethi et al., 2021; Virmani et al., 2021), the extent of hypertrophy seen in this study is most often associated with neurological disorders or normal ageing (Lindsey et al., 1979; Wang et al., 2006; Rodríguez et al., 2014; Joe et al., 2018; Verkhratsky et al., 2019; Bondi et al., 2021). In particular, hypertrophic astrocytes in the hippocampus appear from about 14 months and beyond in ageing rodents (Rodríguez et al., 2014; Matias et al., 2019; Bondi et al., 2021). This hints at the possibility that the process of ageing may be accelerated in MS animals, although this would require a comprehensive investigation. It would be interesting to perform astrocyte-specific transcriptomics studies to understand the nature of these hypertrophic astrocytes and their influence on behavioral deficits. Given the persistent adverse effects of ELS on synaptic function and behavior, it is plausible that these astrocytes may express the genes that are detrimental to synapse survival, as seen with normal ageing (Clarke et al., 2018). This will need a careful examination of the precise age at which hypertrophy starts in ELS-exposed mice and mRNA sequencing during various stages of astrogliosis, since the reactive astroglial transcriptome is highly dynamic and may change over time (Hasel et al., 2021). This study opens up several new avenues to explore the nature of hypertrophic astrocytes in maternally separated mice.

In summary, by performing detailed morphometric analysis, we show that hippocampal astrocytes exhibit significant hypertrophy in maternally separated mice at 8 months of age. This data shows that the effects of ELS on hippocampal astrocytes are long-lasting; and hence, may be associated with the persistent adverse effects of ELS. Further studies are needed to understand the role these hypertrophic astrocytes play in the precipitation of the behavioral impairments following ELS.

## Abbreviations

CNS: Central nervous system
ELS: Early-life stress
GFAP: Glial Fibrillary Acidic Protein
MS: Maternal separation
P2: postnatal day 2
P22: postnatal day 22
PAP: Perisynaptic astrocytic process
PB: Phosphate buffer
PFA: Paraformaldehyde.

## Acknowledgements

Authors would like to thank Mr Manjunath and the staff at the central animal facility and the bioimaging facility at IISc for technical help.

## Author contributions

Conceptualization: S.M.; Methodology: A.N., G.V.; Software: A.N.; Formal analysis: A.N.; Investigation: A.N., G.V., S.M.; Resources: S.M.; Writing - original draft: S.M.; Writing - review & editing: A.N., G.V., S.M.; Supervision: S.M.; Project administration: S.M.; Funding acquisition: S.M.

## Data Availability Statement

The code used for morphometric analysis is available at https://github.com/swanandlab/SMorph. The original images used for analysis and the numerical data can be made available upon reasonable request to the corresponding author.

