## Supplemental file for "Hippocampal astroglial hypertrophy in mice subjected to early life maternal deprivation"

***Figure legends:***

**Figure S1:**

**Analysis of the average lengths of primary, secondary and tertiary processes of hippocampal astrocytes subjected to maternal separation**

We analyzed the average lengths of various branch classes, but found no statistically significant difference in the average length of **(A)** primary, **(B)** secondary and **(C)** tertiary processes between MS animals and controls. n= 90-100 cells per group sampled evenly from 4 mice per group. All comparisons made using Mann-Whitney *U*-test (A,B) or Student's unpaired *t*-test (C).

### FIG S1

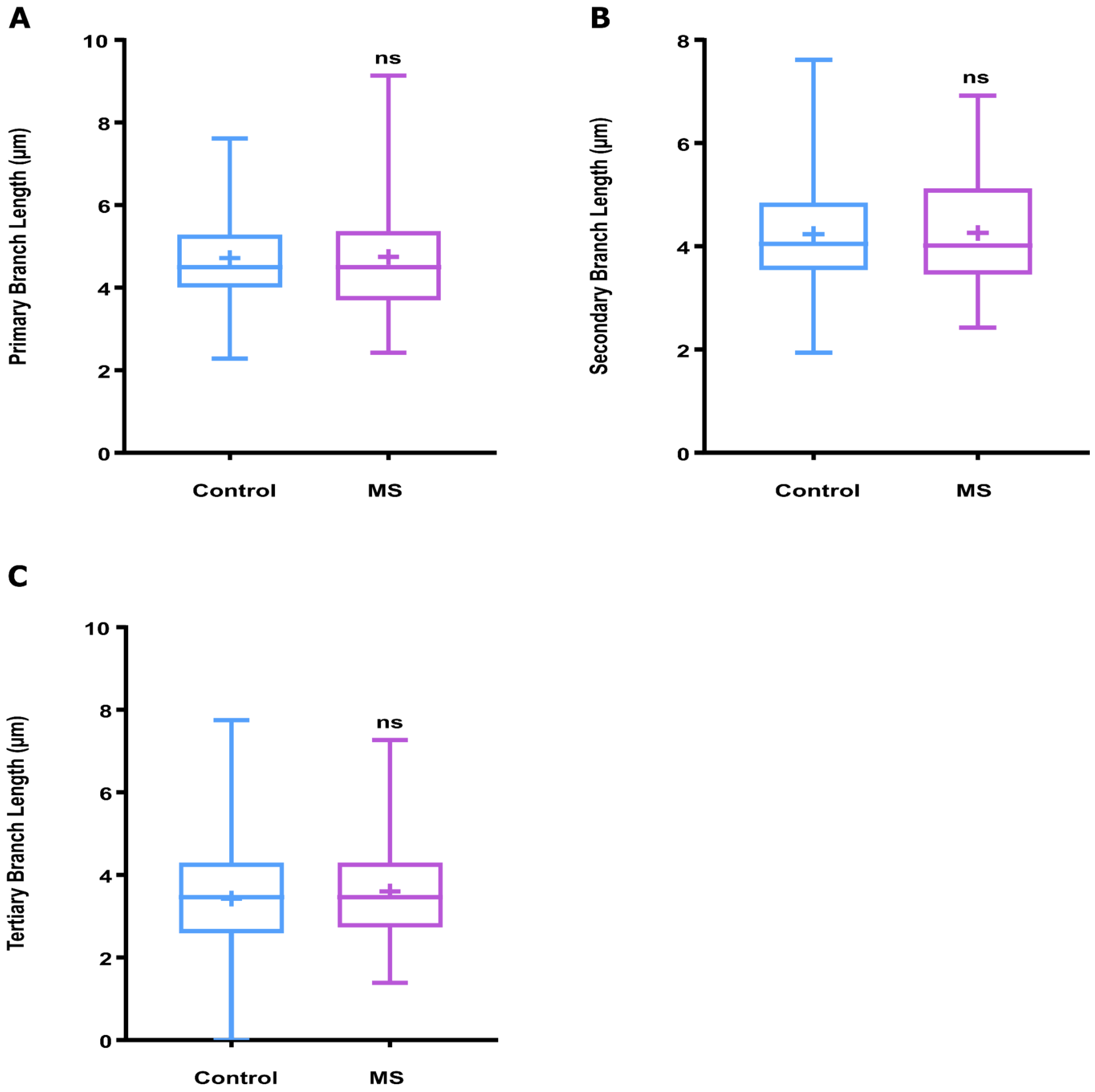
A
10
8
6
4
2
0
Primary Branch Length (µm)
ns
Control
MS
B
8
6
4
2
0
Secondary Branch Length (µm)
ns
Control
MS
C
10
8
6
4
2
0
Tertiary Branch Length (µm)
ns
Control
MS
